# The AGE receptor, OST48 drives podocyte foot process effacement and basement membrane expansion in experimental diabetic kidney disease via promotion of endoplasmic reticulum stress

**DOI:** 10.1101/710186

**Authors:** Aowen Zhuang, Felicia YT Yap, Domenica McCarthy, Sally A. Penfold, Karly C. Sourris, Melinda T. Coughlan, Benjamin L Schulz, Josephine M Forbes

## Abstract

The accumulation of advanced glycation end products is implicated in the development and progression of diabetic kidney disease. No study has examined whether stimulating advanced glycation clearance via receptor manipulation is reno-protective in diabetes. Podocytes, which are early contributors to diabetic kidney disease and could be a target for reno-protection. To examine the effects of increased podocyte oligosaccharyltransferase-48 on kidney function, glomerular sclerosis, tubulointerstitial fibrosis and proteome (PXD011434), we generated a mouse with increased oligosaccharyltransferase-48kDa subunit abundance in podocytes driven by the podocin promoter. Despite increased urinary clearance of advanced glycation end products, we observed a decline in renal function, significant glomerular damage including glomerulosclerosis, collagen IV deposition, glomerular basement membrane thickening and foot process effacement and tubulointerstitial fibrosis. Analysis of isolated glomeruli identified enrichment in proteins associated with collagen deposition, endoplasmic reticulum stress and oxidative stress. Ultra-resolution microscopy of podocytes revealed denudation of foot processes where there was co-localization of oligosaccharyltransferase-48kDa subunit and advanced glycation end-products. These studies indicate that increased podocyte expression of oligosaccharyltransferase-48kDa subunit results in glomerular endoplasmic reticulum stress and a decline in kidney function.

## Introduction

There is a rising global pandemic of diabetes[1, 2], defined by persistent hyperglycemia, a principal risk factor for the development of concomitant chronic complications[3, 4]. Hence, the numbers of individuals with diabetic kidney disease (DKD), a major complication, are burgeoning. DKD is an important risk factor for both end-stage kidney disease and cardiovascular disease[3, 5]. While renin-angiotensin system blockade including angiotensin-converting-enzyme inhibition is first line therapy for DKD, in general these agents only achieve a ∼ 20% reduction in end-stage kidney disease[6].

Advanced glycation end products (AGEs) are a heterogeneous and complex group of non-enzymatic, post-translational modifications to amino acids and proteins, which include hemoglobin A1C (HbA1C) a clinical marker used for the diagnosis of diabetes. Their endogenous formation can be exacerbated by chronic hyperglycemia and oxidative stress [7] and therefore the accumulation of AGEs occurs at an accelerated rate in diabetes[8, 9]. Increases in AGE formation on skin collagen[10-12] and within the circulation[13, 14] predict poor prognosis for patients with diabetes including increased risk for kidney and cardiovascular disease. The kidneys are a major site of AGE detoxification through the filtration of circulating AGEs and their subsequent active uptake and excretion[15, 16]. Therefore, AGE accumulation is seen in diabetic patients with chronic kidney disease[17, 18]. Therapies which lower AGE accumulation have shown benefit in Phase II clinical trials for the treatment DKD in individuals with Type 2 diabetes[19].

OST48, an evolutionarily conserved type 1 transmembrane protein[20], is encoded by the *DDOST* (Dolichyl-diphosphooligosaccharide—protein glycosyltransferase 48 kDa subunit) gene and has been postulated to function as a receptor for AGE turnover and clearance[21, 22]. OST48 is expressed in most cells and tissues[23], including macrophages[24] and mononuclear cells[25]. In the kidney, previous studies have shown OST48 expression in glomerular cells including podocytes[23, 26] and mesangial cells[24] where in the podocytes it is postulated to mediate the uptake and secretion of AGEs[27]. The process whereby AGEs are cleared by OST48 in the kidney is not well understood[28], but is thought to involve degradation of AGEs and then excretion via the urine[29]. A strong link exists between impaired podocyte function and albuminuria, increased urinary AGE excretion and glomerulosclerosis[30-35].

Early damage to the glomeruli, specifically podocyte structural damage, is characteristic of DKD [36-38]. AGEs can induce podocyte cell-cycle arrest[39], hypertrophy[39] and apoptosis[40], while a systemic reduction in AGEs has been shown to improve podocyte and kidney function[41]. Hence, increased facilitation of AGE excretion by increasing OST48 expression could potentially improve podocyte health and alleviate DKD. Currently there are no reported studies identifying whether podocyte OST48 facilitates urinary AGE excretion and subsequently if targeting podocyte OST48 could influence kidney health.

Here, we examined if facilitating greater AGE clearance via modestly increasing podocyte-specific OST48 expression could attenuate the development and progression of DKD.

## Results

### Generation of a podocyte-specific OST48 knock-in mouse model

Mice were generated with a podocyte specific knock-in of human OST48 (*DDOST*+*/-*^Pod-Cre^) driven by the podocin promoter (**Supplementary Information 1A-B**) inserted at the ROSA26 locus and showed no differences in any anthropometric parameters or long-term glycemic control (**Table 1**). We have previously confirmed that this genetic modification does not affect *N*-glycosylation machinery[42]. Mice with experimentally induced diabetes were characterized by elevated fasting blood glucose and glycated hemoglobin (GHb) (**Table 1**). Irrespective of genotype, diabetic mice exhibited a lower body weight, renal hypertrophy, increased consumption of food and water and a greater urinary output when compared with non-diabetic mice (**Table 1**).

**Table 1.**
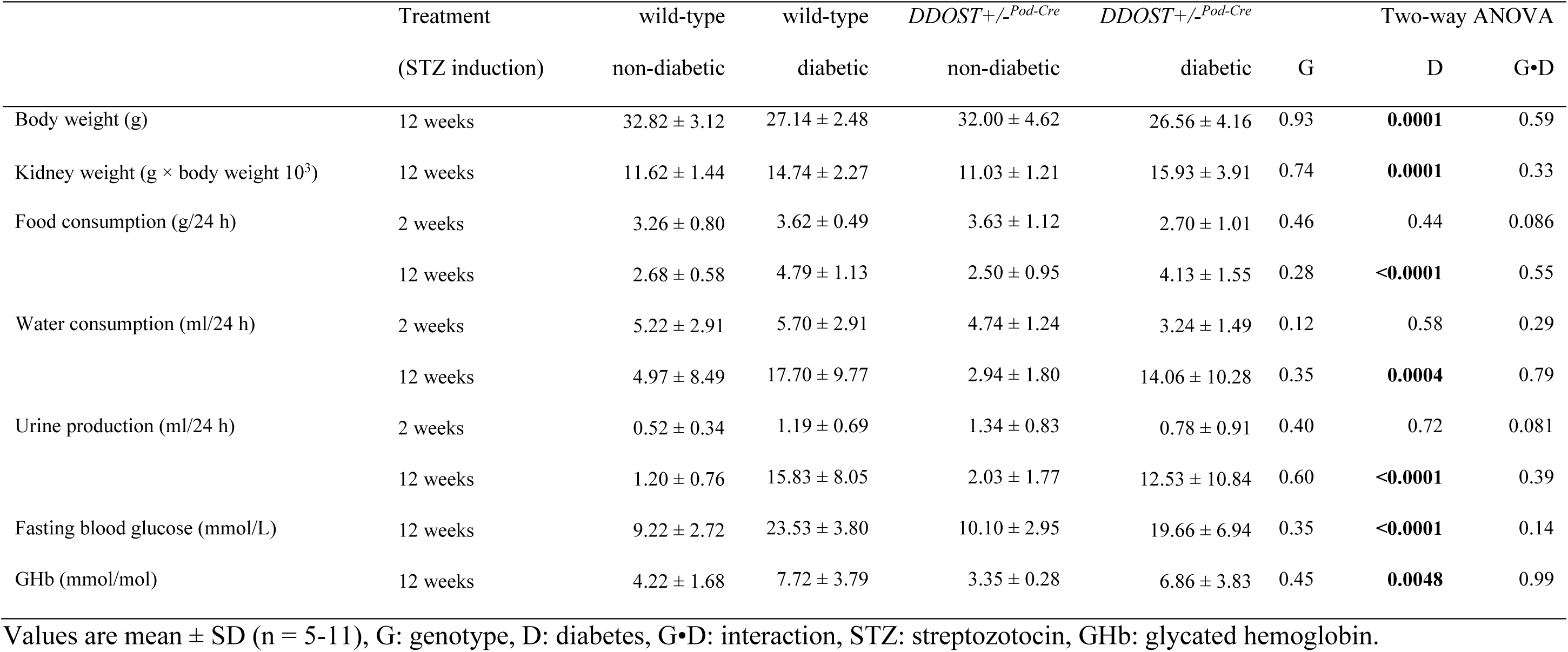
Anthropometric and metabolic parameters.

### *DDOST*+*/-*^Pod-Cre^ mice had podocyte specific increases in OST48 protein content

SWATH-MS proteomics measured the relative abundancy of OST48 in renal cortices (**Figure 1A**) and identified that *DDOST*+*/-*^Pod-Cre^ mice had a significant increase in OST48 protein abundancy (2.06-fold). Moreover, SWATH-MS proteomes of glomeruli (**Figure 1B; left**) or tubular-enriched (**Figure 1B; right**) fractions revealed that *DDOST*+*/-*^Pod-Cre^ mice specifically had significant increases in OST48 abundancy (2.59-fold and 1.79-fold, respectively). Interestingly, it appeared that there was also a significant interactive effect of diabetes on the abundancy of OST48 in glomeruli (**Figure 1B; left**). Localized increases in OST48 abundance in WT-1 positive cells were demonstrated in *DDOST*+*/-*^Pod-Cre^ mice by confocal microscopy (**Figure 1C**). A podocyte-specific increase in OST48 expression was confirmed using ultra resolution 3-dimensional structured illumination microscopy (3D-SIM) (**Figure 1D**). This included increased localization of OST48 to damaged podocyte foot processes, as indicated by nephrin loss (**Figure 1D**), which was particularly evident in 3D reconstruction videos (**Supplementary Figure 2A** and **Supplementary Video 1**). This localization of OST48 to areas of damaged foot processes was unexpected, since the prevailing model is that increasing OST48 abundance could improve declining kidney function by sequestering AGEs[43]. These initial findings of damage to podocyte foot processes warranted further investigation of kidney function in *DDOST*+*/-*^*Pod-Cre*^ mice, and whether increasing podocyte expression of OST48 affected glomerular filtration.

**Figure 1.**
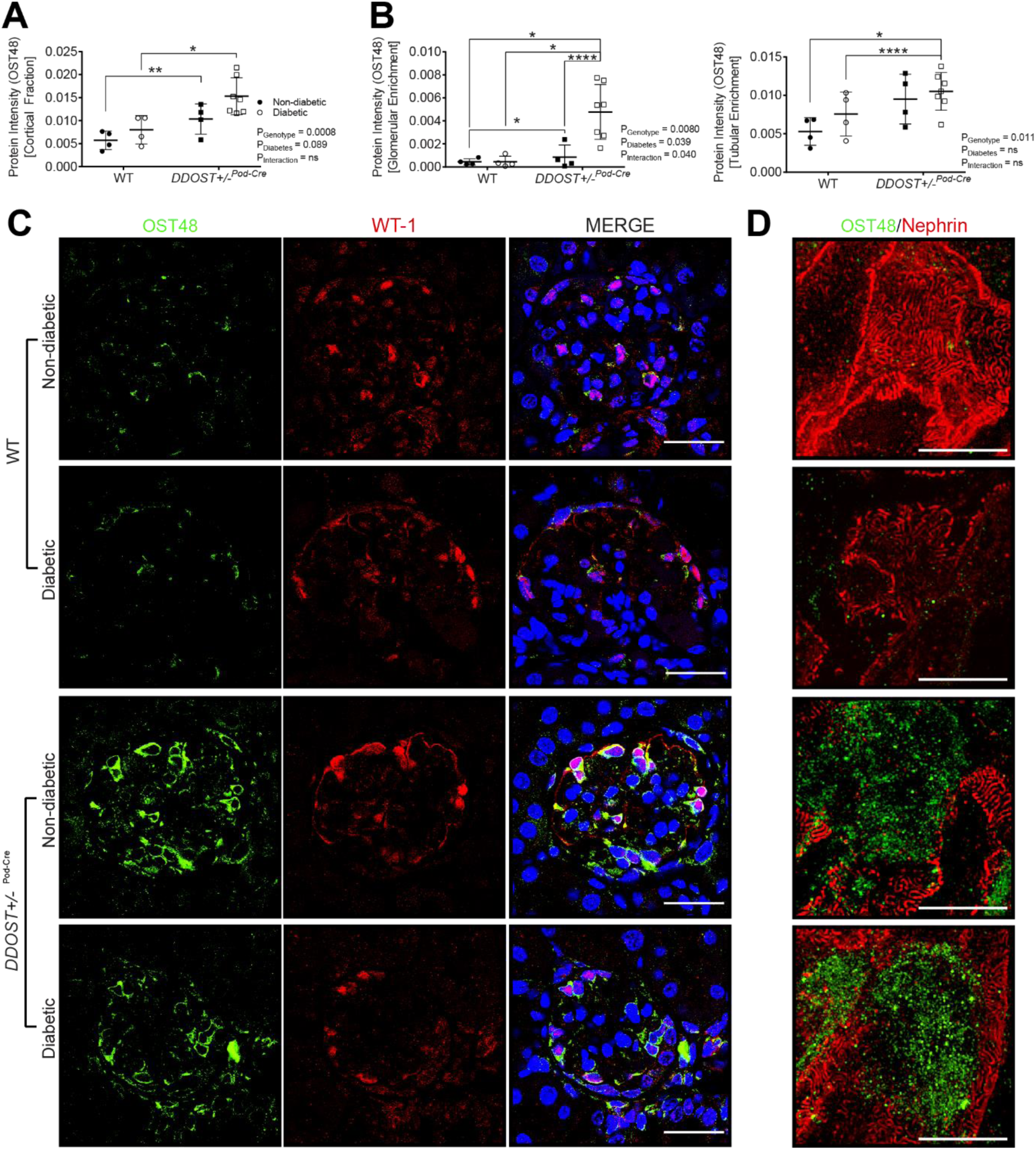
OST48 expression and localization in the kidney at study completion. (**A-B**) SWATH-MS proteomics quantification of total OST48 in (**A**) renal cortex and (**B**) enriched for either (**left**) glomeruli or (**right**) tubules in both the *DDOST*+*/-*^Pod-Cre^ mice and their wild-type littermates. (**C**) Confocal photomicrographs of OST48 (green) and a podocyte specific marker WT-1 (red) on kidney sections imaged at the glomerulus. (**D**) 3D-SIM of podocyte foot processes stained with nephrin (red) and OST48 localization (green). Scale bars from representative images for (**C**) confocal microscopy and (**D**) 3D-SIM were 30µm and 5µm, respectively. Results are expressed as mean ± SD with unpaired t-test analysis (n = 6-12) \*\**P* < 0.01, \*\*\**P* < 0.001 and for proteomics (n = 4-6), MSstatsV2.6.4 determined significant changes in the protein intensities * *P* < 0.05, \*\**P* < 0.01, \*\*\*\**P* < 0.0001.

### Increases in podocyte OST48 expression decreases GFR, exacerbating DKD

Conscious glomerular filtration rate (GFR), determined as a ratio of subcutaneous excretion of FITC-sinistrin, identified a significant decline in kidney function in *DDOST*+*/-*^Pod-Cre^ mice (**Figure 2A** – 49% decrease, *P* < 0.0001). Compared to their baseline kidney function, *DDOST*+*/-*^Pod-Cre^ mice lost 59% of GFR over the study period (**Figure 2B**). Non-diabetic *DDOST*+/-^Pod-Cre^ mice also had elevated serum creatinine compared to their wild-type counterparts (**Figure 2C** – average 88% increase; *P* = 0.0034), which was further increased by diabetes (**Figure 2C** – 58% increase; *P* = 0.043 for *DDOST*+*/-*^Pod-Cre^ and 123% increase; *P* = 0.0047 for wild-type). Diabetes increased serum creatinine in wild type mice (**Figure 2C**) and decreased creatinine clearance (**Figure 2D**), which tended to further decrease in *DDOST*+*/-*^Pod-Cre^ mice, although this did not reach statistical significance (**Figure 2D**). Compared to wild-type mice, *DDOST*+*/-*^Pod-Cre^ mice averaged a 72% reduction in creatinine clearance following simultaneous blood and 24-hour urine collection (**Figure 2E**; *P* = 0.0052) in agreement with FITC-sinistrin based GFR assessment (**Figure 2A**). Diabetes increased 24-hour urinary albumin excretion rate, but this did not differ between wild-type and *DDOST*+*/-*^Pod-Cre^ mice (**Figure 2F and 2G**).

**Figure 2.**
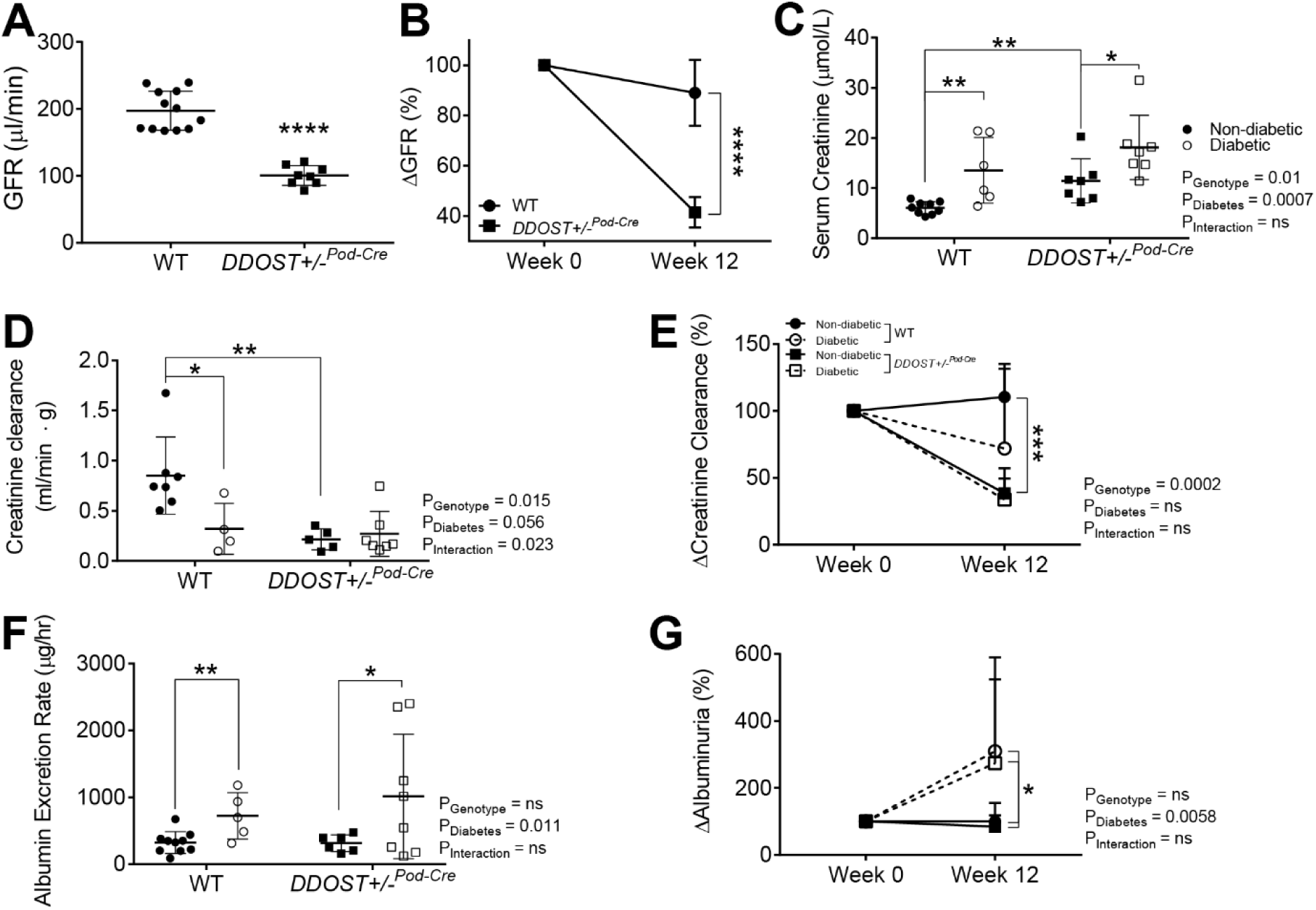
*DDOST*+*/-*^Pod-Cre^ mice exhibited a rapid and progressive decline in kidney function independent of macroalbuminuria. (**A**) Calculated GFR using the transcutaneous decay of retro-orbitally injected FITC-sinistrin. (**B**) Change in GFR, which was determined from matched GFR values from week 0 of the study (6-8 weeks of age) to week 12 of the study. Serum and 24-hour urine was collected from mice in metabolic cages at week 0 and week 12 of the study. (**C-E**) Creatinine was measured spectrophotometrically at 550nm in a biochemical analyzer. (**C**) Serum creatinine measured at week 12 of the study. (**D**) Ratio of creatinine clearance following correction for kidney weight measured at week 12 of the study. (**E**) Change in creatinine clearance, which was determined from matched creatinine clearance values from week 0 of the study (6-8 weeks of age) to week 12 of the study. (**F-G**) Albumin was measured spectrophotometrically at 620nm in a biochemical analyzer and the albumin excretion rate (AER) was determined based on the 24-hour urine flow rate. (**F**) AER measured at week 12 of the study. (**G**) Change in albuminuria over the study, which was determined from matched albuminuria values from week 0 (6-8 weeks of age) of the study to week 12 of the study. Results are expressed as mean ± SD with either two-way ANOVA or paired t-test analysis (n = 3-8) \**P <* 0.05, \*\**P* < 0.01, \*\*\**P* < 0.001.

### *DDOST*+*/-*^Pod-Cre^mice have damaged podocyte architecture and tubulointerstitial damage

Diabetes resulted in glomerulosclerosis in both genotypes (**Figure 3A**). In the absence of diabetes, *DDOST*+*/-*^Pod-Cre^ mice also had glomerulosclerosis (**Figure 3A**) and greater glomerular collagen IV accumulation (**Figure 3B**) than wild-type mice which was not further elevated by diabetes. Glomerular collagen IV accumulation was also present in all diabetic mice (**Figure 3B**). By electron microscopy, glomerular basement membrane thickening and podocyte foot process effacement were seen in *DDOST*+*/-*^Pod-Cre^, but not in wild-type mice (**Figure 3C**). In glomerular fractions from mouse kidney cortices, SWATH-MS proteomics identified significant increases in the abundance of collagen proteins (**Figure 3D**). Specifically, both diabetes and increased podocyte OST48 expression caused an increased abundance of collagen 1, collagen 6 and other structural collagen proteins (**Figure 3D**). Given that all *DDOST*+*/-*^Pod-Cre^ mice exhibited a decline in kidney function, we suspected that in addition to damaged glomeruli there would be other structural evidence of progressive kidney damage in the tubules of these mice. As expected, diabetes increased tubulointerstitial fibrosis in WT mice (**Figure 4A-C**). In addition, there was significant tubulointerstitial fibrosis in all *DDOST*+*/-*^Pod-Cre^ mice compared to wild-type non-diabetic mice, evidenced by an increase in positive connective tissue quantified from Masson’s trichrome (**Figure 4A**), Sirius red (**Figure 4B**) and collagen IV staining in this compartment (**Figure 4C**). In tubule-enriched cortical fractions, SWATH-MS proteomics identified modest increases in proteins associated with collagen fibril organization (CO1A1-2) (**Figure 4D**) in non-diabetic *DDOST*+*/-*^Pod-Cre^ mice compared with WT littermates. However, with diabetes, *DDOST*+*/-*^Pod-Cre^ mice showed a significant accumulation of collagen IV in tubule-enriched fractions not seen in WT diabetic or non-diabetic *DDOST*+*/-*^Pod-Cre^ mice. There were further modest increases in markers of tubulointerstitial fibrosis in diabetic *DDOST*+*/-*^Pod-Cre^ but these were not different to WT mice with diabetes (**Figure 4A-D**)

**Figure 3.**
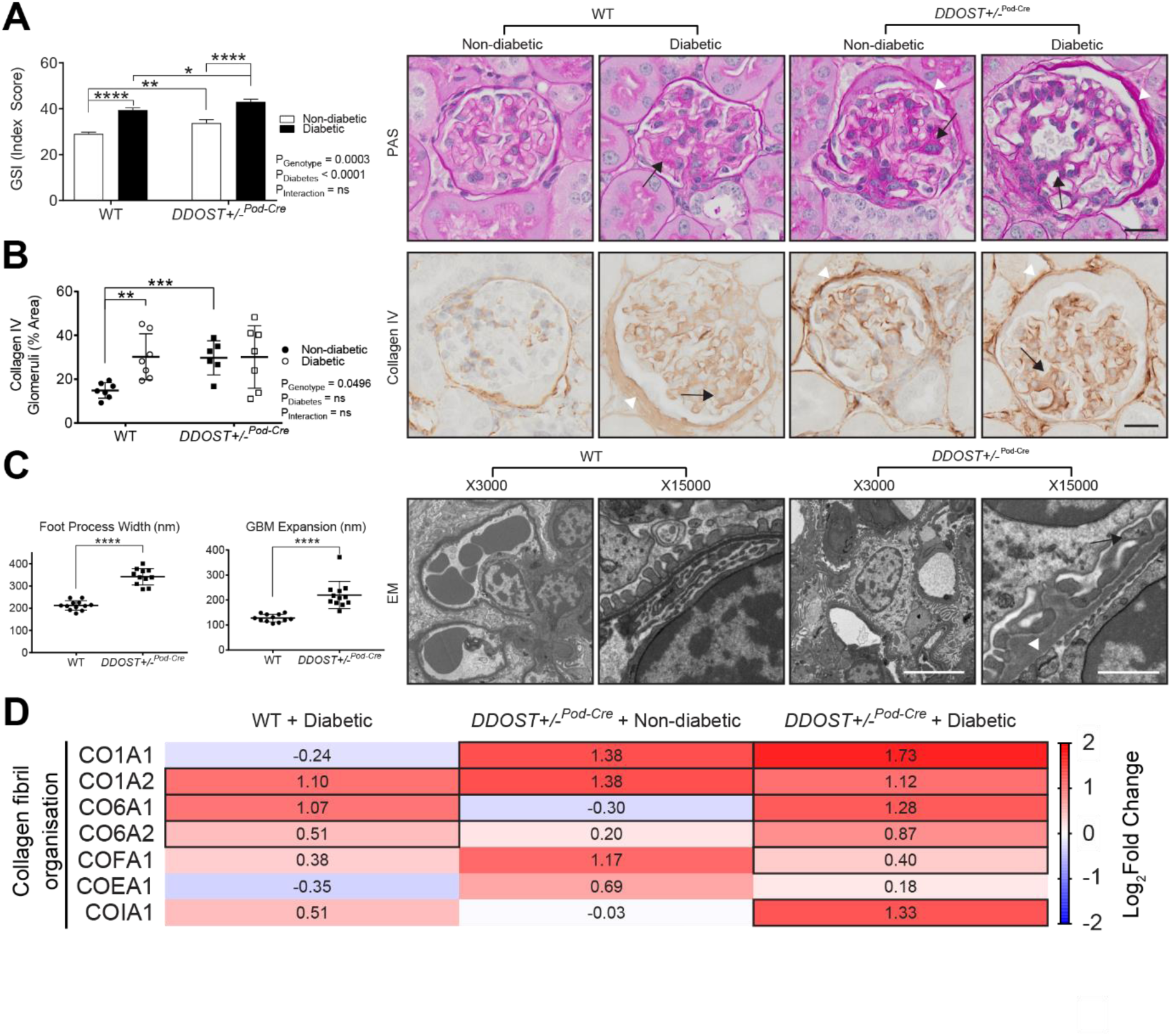
Podocyte OST48 was associated with a deterioration of podocyte architecture. (**A-C**) Assessment of renal glomerular damage in the kidney with (**A**) Periodic-acid Schiff staining (PAS) and (**B**) collagen IV, which was then quantified based on a positive threshold protocol. *DDOST*+*/-*^Pod-Cre^ mice and mice with a diabetic phenotype show moderate glomerulosclerosis, indicated by an increase in mesangial matrix expansion (black arrow) and accumulation of the extracellular matrix (white arrow head). (**C**) Electron microscopy determined the degree of hypertrophy in the foot processes (black arrow) and the expansion of the GBM (white arrow head). Scale bars from representative images of (**A-B**) glomeruli stained with either PAS or collagen IV were 20µm. (**C**) Representative images from electron microscopy were imaged at either X3000 magnification or X15000 magnification were 20µm and 4µm, respectively. (**D**) Heat map representation of SWATH-MS proteomics data for enzymatic pathways involved in collagen fibril organization. Significant proteins are represented as bolded cells, where red indicates an increase and blue indicates a decrease in protein concentrations. Data represented as means ± SD (*n* = 4-7/group). Results are expressed as mean ± SD with either two-way ANOVA or unpaired t-test analysis (n = 5-9) \**P <* 0.05, \*\**P* < 0.01, \*\*\**P* < 0.001, \*\*\*\**P* < 0.0001. For proteomics, MSstatsV2.6.4 determined significant (*P* < 0.05) log fold changes in the protein intensities between the selected experimental group the wild-type non-diabetic group. Heatmap representation allow for compact visualization of complex data comparisons. Full detail of the quantitative data is available in the supplementary information.

**Figure 4.**
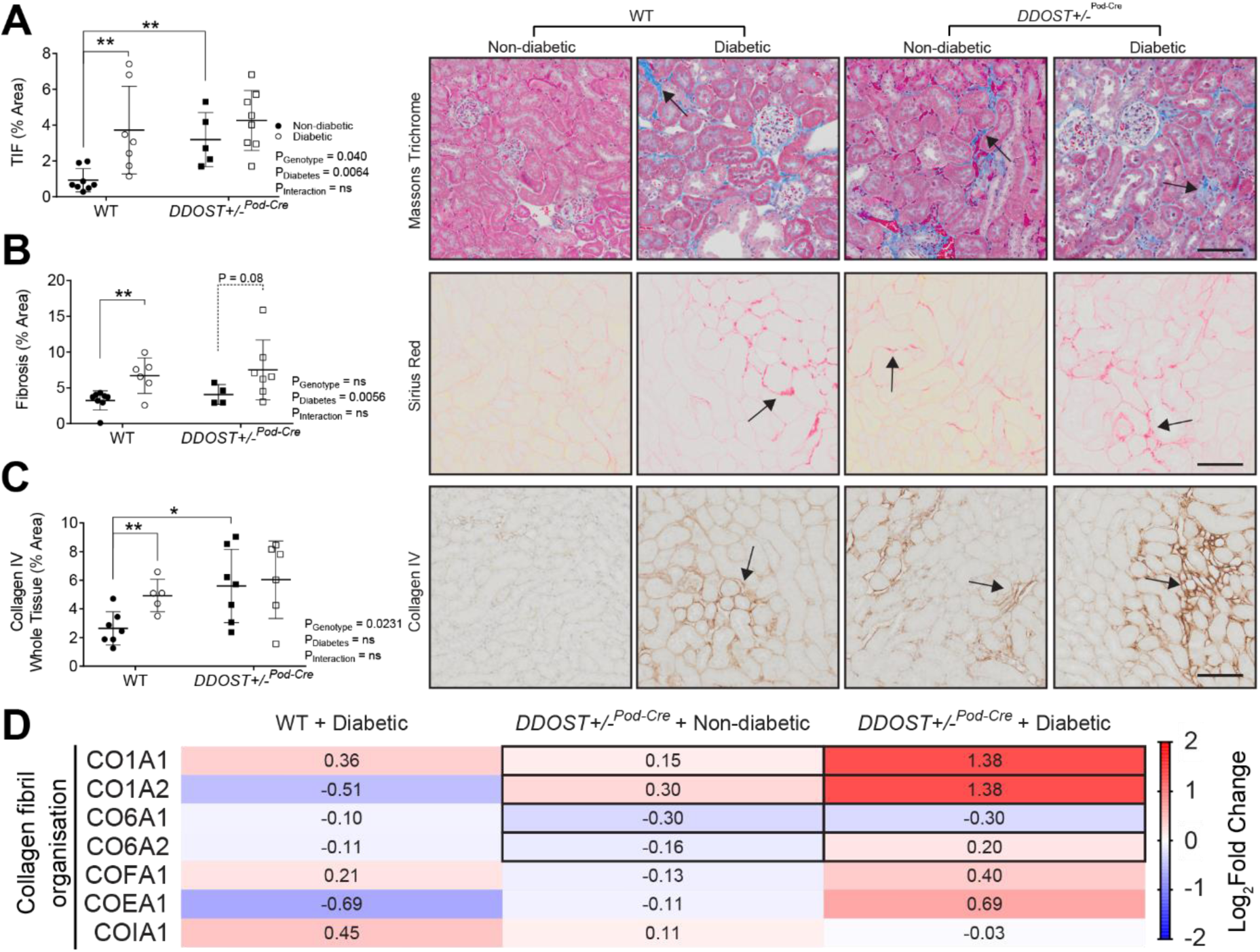
Podocyte OST48 increased tubulointerstitial fibrosis. (**A-C**) Moreover, the presence of collagen in (**A**) Masson’s trichrome (blue staining), (**B**) Sirius red (red) and (**C**) collagen IV (brown), was quantified based on a positive threshold protocol. Fibrosis in the interstitium of the proximal tubules (black arrow) is an indicator of progressive kidney damage. The severity of these changes was more pronounced in the *DDOST*+*/-*^Pod-Cre^ mice and the mice with a diabetic phenotype. (**D**) Heat map representation of SWATH-MS proteomics data for enzymatic pathways involved in collagen fibril organization. Significant proteins are represented as bolded cells, where red indicates an increase and blue indicates a decrease in protein concentrations. Data represented as means ± SD (n = 4-7/group). Scale bars from representative images of (**A**-**C**) proximal tubule sections stained with either Masson’s trichrome, Sirius red or collagen IV were 100µm. Results are expressed as mean ± SD with either two-way ANOVA or unpaired t-test analysis (n = 5-9) \**P <* 0.05, \*\**P* < 0.01. For proteomics, MSstatsV2.6.4 determined significant (*P* < 0.05) log fold changes in the protein intensities between the selected experimental group the wild-type non-diabetic group. Heatmap representation allow for compact visualization of complex data comparisons. Full detail of the quantitative data is available in the supplementary information.

### Tissue AGEs were localized to damaged podocytes

Podocyte AGE accumulation and OST48 appeared to be co-localized (**Figure 5A**) and were prominent in areas where concomitant deterioration of podocyte foot processes was seen, as defined by loss of both nephrin (**Figure 5A**) and synaptopodin (**Supplementary Figure 3A**). 3D-SIM confirmed the appearance of nodules/vacuoles containing both OST48 and AGEs in *DDOST*+*/-*^Pod-Cre^ mice in areas of podocyte foot process deterioration (**Figure 5B, Figure 5C** and **Supplementary Video 2**). These changes were not observed in wild-type diabetic mice. Overall, there was a significant reduction in AGE content in kidney cortical extracts taken from *DDOST*+*/-*^Pod-Cre^ mice (**Figure 5D** - genotype effect *P* < 0.05). Diabetes also increased urinary AGE excretion (*P* = 0.007; ∼10-fold increase) when compared to non-diabetic mice (**Figure 5E**). Although not significant (*P* < 0.2) there was also a trend towards increased urinary AGE excretion in non-diabetic *DDOST*+*/-*^Pod-Cre^ mice compared to wild-type non-diabetic mice.

**Figure 5.**
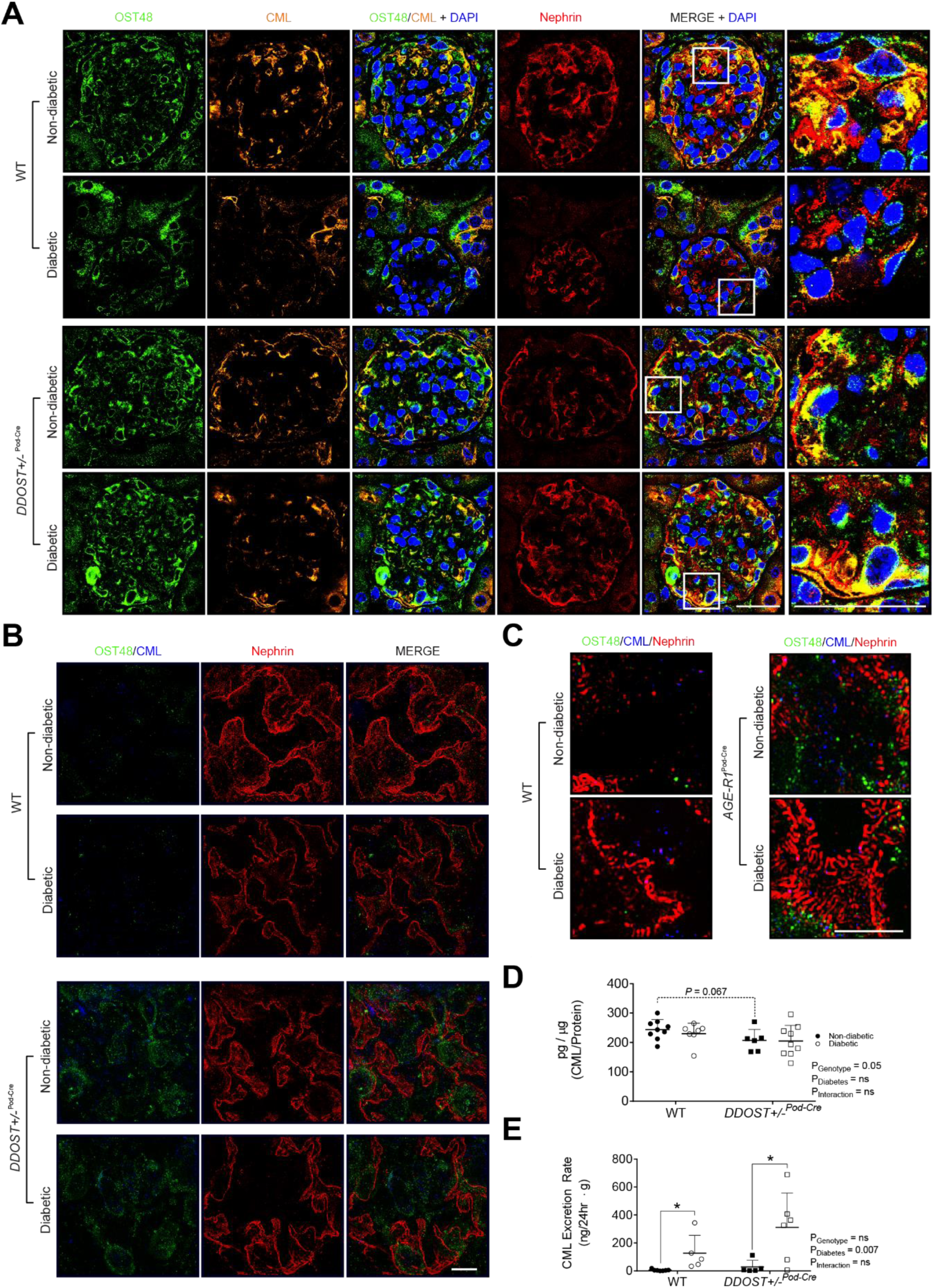
Podocyte OST48 increased AGE accumulation in podocytes. (**A**) Confocal photomicrographs of OST48 (green), CML (orange) and a podocyte foot process marker, nephrin (red) on kidney sections imaged at a glomerulus. (**B**) 3D-SIM of podocyte foot processes staining for nephrin (red), OST48 (green) and CML (blue) localization. (**C**) 3D-SIM reconstruction of podocyte foot processes staining for nephrin (red), OST48 (green) and CML (blue) localization. (**D-E**) CML ELISA measuring the total content of CML detected in (**D**) whole kidney cortex homogenates and in (**E**) 24-hour urine collections. Scale bars from representative images for (**A**) confocal microscopy and (**B-C**) 3D-SIM were 30µm and 5µm, respectively. Results are expressed as mean ± SD with either two-way ANOVA or unpaired t-test analysis (n = 5-9) \**P <* 0.05.

### Podocyte OST48-mediated AGE accumulation resulted in ER stress

ER stress at sites of AGE accumulation within damaged podocytes was examined. *DDOST*+*/-*^Pod-Cre^ mice exhibited an increase in the ER stress markers GRP-78 and spliced XBP-1 when compared to wild-type mice (**Figure 6A**). Furthermore, these ER stress markers were further increased by diabetes (**Figure 6A**). This was confirmed using SWATH-MS proteomics of glomerular enriched fractions, where diabetic *DDOST*+*/-*^Pod-Cre^ mice showed significantly higher abundance of proteins associated with ER stress (GRP75 and GRP78) compared to non-diabetic wild-type mice (**Figure 6B**). Antioxidant enzymes SOD1/SODC and SOD2/SODM were also increased by diabetes in wild-type mice (**Figure 6B**). Compared with non-diabetic mice, glomerular fractions from diabetic wild-type and *DDOST*+*/-*^Pod-Cre^ mice had significantly higher levels of proteins associated with mitochondrial ATP synthesis, particularly biosynthesis of complex I components (NADH dehydrogenase:ubiquinone) (**Figure 6C**).

**Figure 6.**
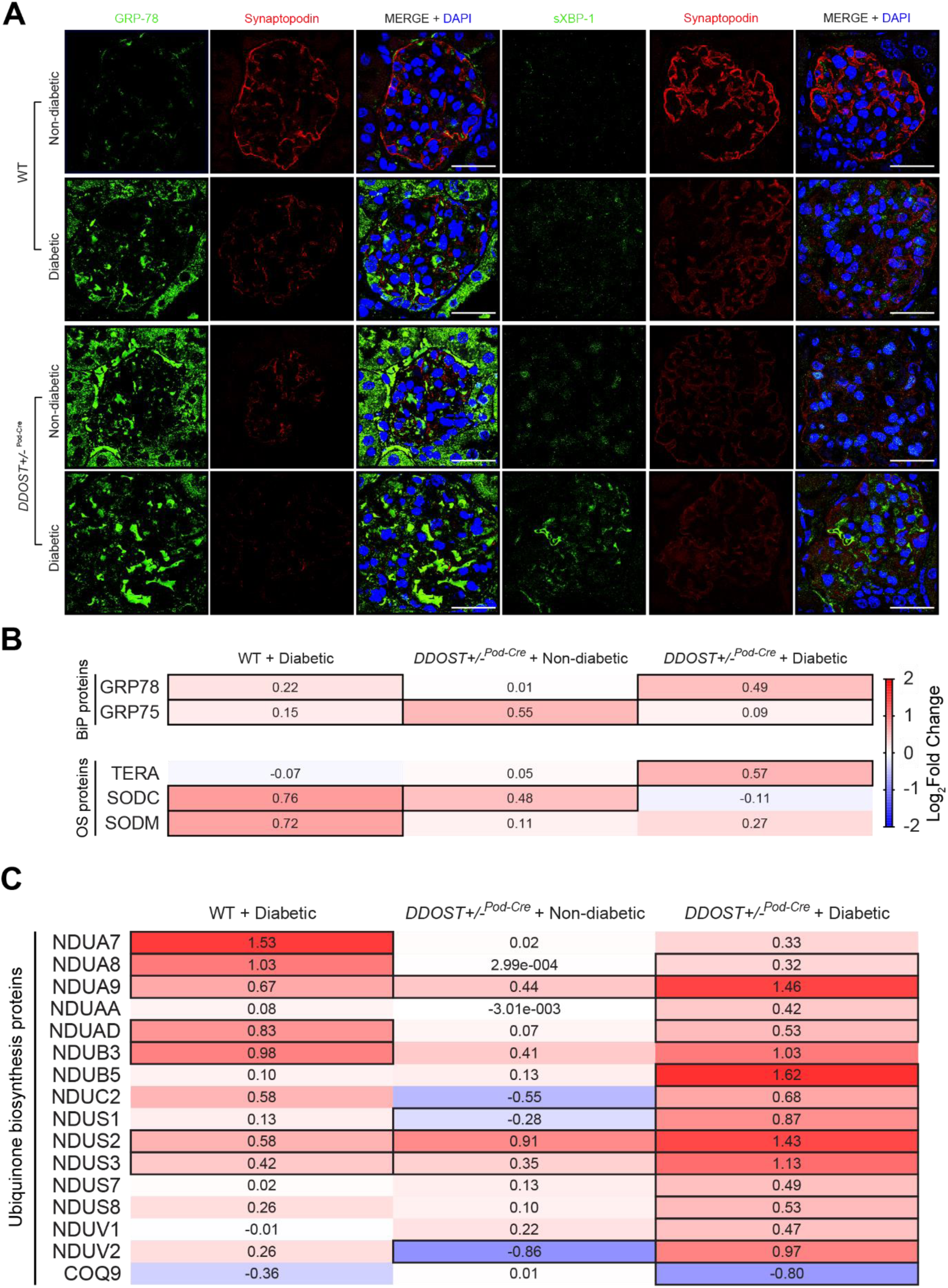
Podocyte OST48 increased ER stress markers. (**A**) Confocal photomicrographs of either ER stress marker GRP-78 (green), or XBP-1 (green) and podocyte foot process marker, synaptopodin (red). (**B**-**C**) Heat map representation of SWATH-MS proteomics data for enzymatic pathways involved in (**B**) endoplasmic reticulum (ER) stress and oxidative stress (OS) enzymatic pathways, and (**C**) ubiquinone biosynthesis pathways. Significant proteins are represented as bolded cells, where red indicates an increase and blue indicates a decrease in protein concentrations. Data represented as means ± SD (n = 4-7/group). Scale bars from representative images of confocal microscopy were 30µm. Data represented as means ± SD (n = 5/group). For proteomics, MSstatsV2.6.4 determined significant (P < 0.05) log fold changes in the protein intensities between the selected experimental group the wild-type non-diabetic group. Heatmap representation allow for compact visualization of complex data comparisons. Full detail of the quantitative data is available in the supplementary information.

## Discussion

Lowering AGE burden, including by facilitating renal AGE clearance, has been suggested as a possible treatment for DKD. However, the assertion that increasing renal AGE clearance receptors such as OST48 may protect against DKD has not been previously investigated. Here, we have shown for the first time that increasing OST48 in a podocyte-specific manner decreased glomerular filtration and caused podocyte structural damage including foot process effacement, leading to glomerulosclerosis and tubulointerstitial fibrosis, which were further exacerbated by diabetes. Surprisingly, in non-diabetic mice the decline in GFR and increased podocyte and glomerular damage were seen in the absence of elevations in albuminuria. Mechanistically, increases in OST48 in podocytes were marked by increased podocyte specific AGE uptake resulting in oxidative and endoplasmic reticulum stress, which is independent of changes to the *N*-glycosylation machinery and subsequent kidney functional and structural decline (**Figure 7**), where previously it has been shown that AGE uptake into cultured podocytes induces hypertrophy[39] and apoptosis[40].

**Figure 7.**
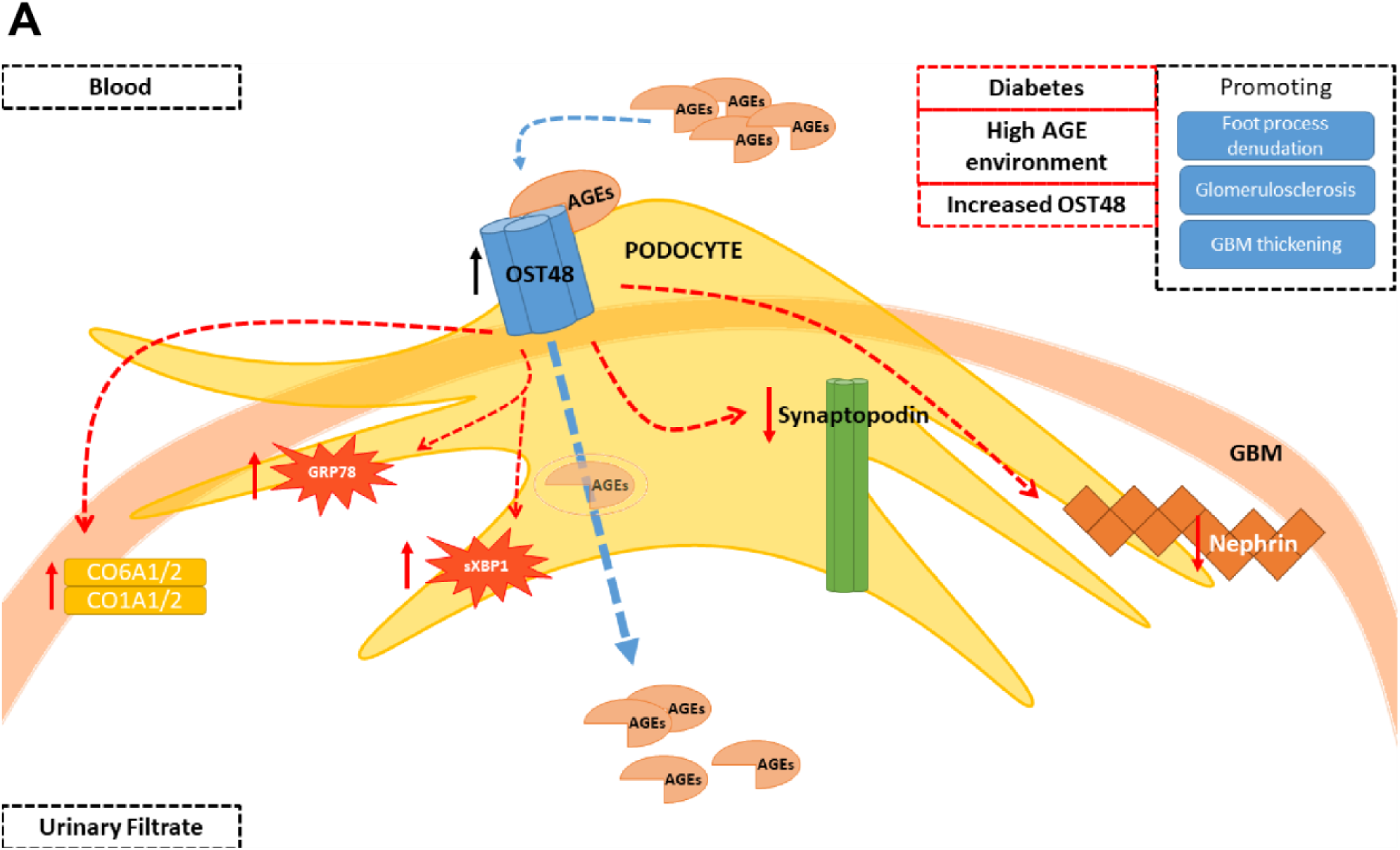
Podocyte OST48 drives podocyte foot process effacement. (**A**) Working diagram depicting the model whereby OST48 drives podocyte foot process effacement in a streptozotocin-induced model of DKD. AGEs; advanced glycation end products, GBM; glomerular basement membrane, OST48; advanced glycation end-product receptor 1/oligosaccharyltransferase-48kDa subunit, GRP78; 78 kDa glucose-regulated protein/binding immunoglobulin protein, sXBP1; spliced X-box binding protein 1; CO6A1/2; collagen alpha-1/2 (VI) chain; CO1A1/2; collagen alpha-1/2 (I) chain.

It was particularly interesting that albuminuria was not seen concomitantly with the loss of GFR in non-diabetic *DDOST*+*/-*^*Pod-Cre*^ mice despite podocyte foot effacement, thickening of the glomerular basement membrane and glomerulosclerosis. Although surprising, this is consistent with clinical observations. For example, the prospective longitudinal Joslin Proteinuria Cohort studies[44, 45] identified that the overall prevalence of normo-albuminuria in patients with progressive kidney disease was around 10% and that regression from micro- and macro-albuminuria to normo-albuminuria was repeatedly observed. We observed podocyte foot effacement and glomerular basement membrane thickening, which are commonly attributed as causes of micro- and macro-albuminuria. This finding is consistent with other postulated models of the origins of albuminuria, such as the high glomerular sieving coefficient (GSC) hypothesis[46, 47]. This hypothesis suggests that albuminuria is driven by proximal tubule damage which prevents the reabsorption/processing of albumin by these cells. In support of this model, tubulointerstitial fibrosis was more pronounced in diabetic mice regardless of their genotype, and they also had concomitant increases in urinary albumin excretion which not seen in non-diabetic mice. Alternatively, the absence of macro-albuminuria seen in mice with increased podocyte expression of OST48 could be attributed to the significant decline in GFR, as this would limit the flux of albumin into the urine.

Although we showed that increases in podocyte OST48 expression decreased renal AGE accumulation and increased urinary AGE excretion, this was not associated with reno-protection. In fact, this resulted in significant renal disease. We have shown for the first time that OST48 and AGEs colocalize within podocytes in areas of foot process denudation and damage. We observed that OST48 facilitation of podocyte AGE accumulation resulted in a cascade of ER stress culminating in podocyte foot process effacement, GBM expansion, and renal functional decline. Increasing the urinary flux of AGEs has however been previously shown to impair renal function in both healthy humans[41, 48] and early in the development of diabetic kidney disease in rodent models[35, 49].

Taken together, these results suggest that specifically increasing podocyte exposure and uptake of AGEs by facilitating their flux into the urine is sufficient to induce significant kidney damage.

### Translational Statement

Advanced glycation end-products (AGEs) have long been considered an important pathological factor contributing to diabetic kidney disease (DKD). However, targeting AGEs has proven difficult due to the lack of knowledge in the signaling pathways surrounding its lesser known receptors such as OST48, and how these impact kidney function in health and disease. This paper describes the role of OST48 in the podocytes using genetic manipulation of animal models, proteomics and 3D-SIM microscopy. It demonstrates that facilitation of AGE excretion mediated by OST48 in the podocyte is pathological and detrimental to kidney function through the promotion of podocyte effacement. These results would suggest a paradigm shift in the role currently thought of OST48. This knowledge is key for the development of new therapies targeting advanced glycation or re-purposing known compounds which could provide effective targeting of advanced glycation.

## Methods

### Animals

All animal studies were performed in accordance with guidelines provided by the AMREP (Alfred Medical Research and Education Precinct) Animal Ethics Committee, University of Queensland and the National Health and Medical Research Council of Australia (E/1184/2012/B and MRI-UQ/TRI/256/15/KHA). Male C57BL/6J wild-type mice with a genetic insertion of the human gene encoding AGE-R1/OST-48 (*DDOST*) in the ROSA26 locus (*ROSA26*^*tm1*(*DDOST*)*Jfo*^) crossed with mice only expressing *Cre* under the control of the *Podocin* promoter (termed *DDOST*+*/-*^Pod-Cre^) were generated (**Supplementary Figure 1A**) by Ozgene Australia (Ozgene, Perth, WA, Australia). Between 6-8 weeks of age (Week 0), experimental diabetes (DIAB) was induced in male wild-type (n = 9) and *DDOST*+*/-*^Pod-Cre^ (n = 10) mice by low dose intraperitoneal injection of streptozotocin[50]. Non-diabetic male wild-type (n = 11) and *DDOST*+*/-*^Pod-Cre^ (n = 7) mice received equivalent injections of sodium citrate buffer alone. After 10 days of recovery, blood glucose concentrations were determined and then repeated weekly to ensure mice had diabetes ([blood glucose]>15mmol/L). Mice were allowed access to food and water *ad libitum* and were maintained on a 12-hour light:dark cycle at 22°C. All mice received a diet of standard mouse chow (AIN-93G) (Specialty Feeds, Glen Forrest, Perth, Australia). At 0 and 12 weeks of the study, mice were housed in metabolic cages (Iffa Credo, France) for 24 hours to determine food and water consumption and collect urine. Urine output was also measured during caging and urinary glucose determined by a glucometer (SensoCard Plus, POCD, Australia).

### Glomerular and tubule enrichment

Kidneys were decapsulated and the medulla discarded. Cortex was kept cold, minced, and passed gently through three stacks of fine-mesh sieves (250µm, 70µm and 45µm). Tubular-enriched proteins were contained in the elute whilst the glomerular-enriched proteins were located on the top of the 70µm sieve. Enriched proteins were collected and processed for mass spectrometry analysis. Glomerular-(SYNPO, ACTN4) and tubular-specific (AQP1, DIC, S22A6) proteins were significantly enriched in respective fractions (**Supplementary Figure 1C**).

### Biochemical analyses

Nε-(Carboxymethyl)lysine (CML) concentrations in plasma and urine were measured by an in-house indirect ELISA as previously described[28]. Urinary albumin (Bethyl Laboratories, United States) was normalized to the 24-hour flow rate of urine. Serum and urinary creatinine were measured spectrophotometrically at 505nm (Roche/Hitachi 902 Analyzer, Roche Diagnostics GmBH, Germany) using the Jaffe method.

### Glomerular Filtration Rate

At 0 and 12 weeks of the study, GFR was estimated in conscious mice using the transcutaneous decay of retro-orbitally injected FITC-sinistrin (10mg/100g body weight dissolved in 0.9% NaCl), as previously described[51]. GFR was calculated from the rate constant (α2) of the single exponential excretion phase of the curve and a semi-empirical factor.

### Liquid chromatography-tandem mass spectrometry (LC-MS/MS) proteomics

Fractions enriched for either glomeruli or tubules were processed as previously described[52]. Peptides were desalted and analyzed by Information Dependent Acquisition LC-MS/MS as described[53] using a Prominence nanoLC system (Shimadzu, Australia) and Triple TOF 5600 mass spectrometer with a Nanospray III interface (AB SCIEX, United States). SWATH-MS analysis was performed[54] and differentially abundant proteins were analyzed using DAVID[55]. Processed proteomic data sets are available in the Supplementary Information.

### Histology and imaging

Paraffin-embedded sections were stained with either a Periodic acid Schiff (PAS) staining kit (Sigma-Aldrich, United States), a Trichrome (Masson) staining kit (Sigma-Aldrich, St. Louis, Missouri, USA) or Sirius Red (Sigma-Aldrich, United States). Collagen IV was determined by immunohistochemistry using an anti-collagen IV antibody (1:100 dilution; ab6586; Abcam, United Kingdom). All sections were visualized on an Olympus Slide scanner VS120 (Olympus, Japan) and viewed in the supplied program (OlyVIA Build 10555, Olympus, Japan). Slides were quantified based on threshold analysis in Fiji[56]. Briefly, for immunofluorescence staining, frozen kidney sections were stained with a combination of either anti-CML (1:200 dilution; ab27684; Abcam, United Kingdom), OST48 (H-1; 1:100 dilution; SC-74408; Santa Cruz biotechnologies, United States), GRP78 (A-10; 1:50 dilution; SC-376768; Santa Cruz biotechnologies, United States), WT-1 (c-terminus; 1:100 dilution; PA5-16879; ThermoFisher Scientific, United States), synaptopodin (P-19; 1:100 dilution; SC-21537; Santa Cruz biotechnologies, United States), nephrin (1:100 dilution; ab27684; Abcam, United Kingdom). Confocal images were visualized on an Olympus FV1200 confocal microscope (Olympus, Japan) and viewed in the supplied program (FV10, Olympus, Japan). For ultra-resolution 3D microscopy, slides were visualized on an OMX Blaze deconvolution structured illumination (SIM) super-resolution microscope (GE Healthcare Life Science, United Kingdom).

### Glomerulosclerotic index (GSI)

GSI as a measure of glomerular fibrosis was evaluated in a blinded manner by a semi-quantitative method[57]. Severity of glomerular damage was assessed on the following parameters; mesangial matrix expansion and/or hyalinosis of focal adhesions, true glomerular tuft occlusion, sclerosis and capillary dilation.

### Fixation, tissue processing and acquisition of data from electron microscopy

Renal cortex wa**s** processed as previously described[58]. Sections were cut at 60nm on a Leica UC6 ultramicrotome (Leica, Germany) and were imaged at 80kV on a Jeol JSM 1011 transmission electron microscope (Jeol, United States) equipped with an Olympus Morada (Olympus, Japan) digital camera. Quantification of foot process width and GBM expansion was assessed as previously described[59].

### Statistics

Results are expressed as mean ± SD (standard deviation) and assessed in GraphPad Prism V7.01 for Windows (GraphPad Software, United States). Normally distributed parameters (tested with D’Agostino & Pearson omnibus normality test) were tested for statistical significance by 2-way ANOVA followed by post hoc testing for multiple comparisons using the Bonferroni method unless otherwise specified. For comparison between groups as required, a two-tailed unpaired Student’s t-test was used where specified. For SWATH-MS, MSstatsV2.6.4[60] was used to detect differentially abundant proteins estimating the log-fold changes between compared conditions of the chosen experimental group and with the wild-type non-diabetic mice. For all calculations, a *P* < 0.05 was considered as statistically significant.

### Data availability statement

The datasets generated and analyzed during the current study are available from the corresponding author upon reasonable request. The mass spectrometry proteomics data have been deposited to the ProteomeXchange Consortium via the PRIDE[61] partner repository with the dataset identifier PXD011434.

## Disclosure

J.M.F. is supported by a Senior Research Fellowship (APP1004503;1102935) from the National Health and Medical Research Council of Australia (NHMRC). B.L.S. is supported by a Career Development Fellowship (APP1087975) from the NHMRC. A.Z. received a scholarship from Kidney Health Australia (SCH17; 141516) and the Mater Research Foundation. M.T.C. is supported by a Career Development Fellowship from the Australian Type 1 Diabetes Clinical Research Network, a special initiative of the Australian Research Council. This research was supported by the NHMRC, Kidney Health Australia and the Mater Foundation, which had no role in the study design, data collection and analysis, decision to publish, or preparation or the manuscript.

## Acknowledgements

The authors acknowledge the Translational Research Institute (TRI) for providing the excellent research environment and core facilities that enabled this research. We particularly thank Sandrine Roy and Ali Ju from the TRI Microscopy Core Facility. Furthermore, the authors acknowledge the assistance in electron microscopy provided by Jennifer Brown from the Centre for Microscopy and Microanalysis at the University of Queensland.

## Author Contributions

A.Z., B.L.S. and J.M.F. designed the study; A.Z., F.Y.T, M.T.C., K.C., D.M. and S.A.P. carried out the experiments; A.Z., B.L.S. and J.M.F. analyzed the data; A.Z., B.L.S. and J.M.F. made the figures; A.Z., B.L.S. and J.M.F. drafted and revised the paper; D.M., K.C.S. and M.T.C. assisted with revision of the paper; all authors approved the final version of the manuscript.

