## Supplementary for "The AGE receptor, OST48 drives podocyte foot process effacement and basement membrane expansion in experimental diabetic kidney disease via promotion of endoplasmic reticulum stress"

**Supplementary Figure Legends**

**Supplementary Figure 1.** Generation of a podocyte specific *DDOST* heterozygous knock-in mutant. (**A**) Genomic clone of DDOST bearing exon 1 used for the construction of the floxed targeting vector as indicated. Heterozygous mice were crossed with mice expressing podocin-Cre and subsequent progeny contained a ubiquitous over-expression allele. (**B**) Genomic Southern blotting confirmed the predicted DDOST allelic structures in the podocin-targeted knock-in strain. (**C**) Protein intensities of SWATH-MS proteomics data for podocyte specific proteins (sp|Q8CC35|SYNPO_MOUSE and sp|P57780|ACTN4_MOUSE) and proximal tubule specific proteins (sp|Q02013|AQP1_MOUSE, sp|Q9QZD8|DIC_MOUSE and sp|Q8VC69|S22A6_MOUSE) enriched in the glomerular or tubule fractions.

**Supplementary Figure 2.** (**A**) Reconstruction of 3D-SIM of podocyte foot processes stained with nephrin (red) and OST48 localization (green). Scale bars from representative images for 3D-SIM were 5µm.

**Supplementary Figure 3.** Podocyte OST48 increased AGE accumulation in podocytes. (**A**) Confocal photomicrographs of OST48 (green), CML (orange) and a podocyte foot process marker, synaptopodin (red) on kidney sections imaged at a glomerulus. Scale bars from representative images for confocal microscopy were 30µm. (**B**) ELISA measuring the total content of CML detected in plasma.

**Supplementary Video 1.** Reconstruction of 7µm thick renal cortex section imaged in a 3D-SIM microscope. OST48 (green), nephrin (red). Scale bars from representative videos for 3D-SIM were 5µm.

**Supplementary Video 2.** Reconstruction of 7µm thick renal cortex section imaged in a 3D-SIM microscope. OST48 (green), CML(blue) and nephrin (red). Scale bars from representative videos for 3D-SIM were 5µm.

### Supplementary Figure 1.

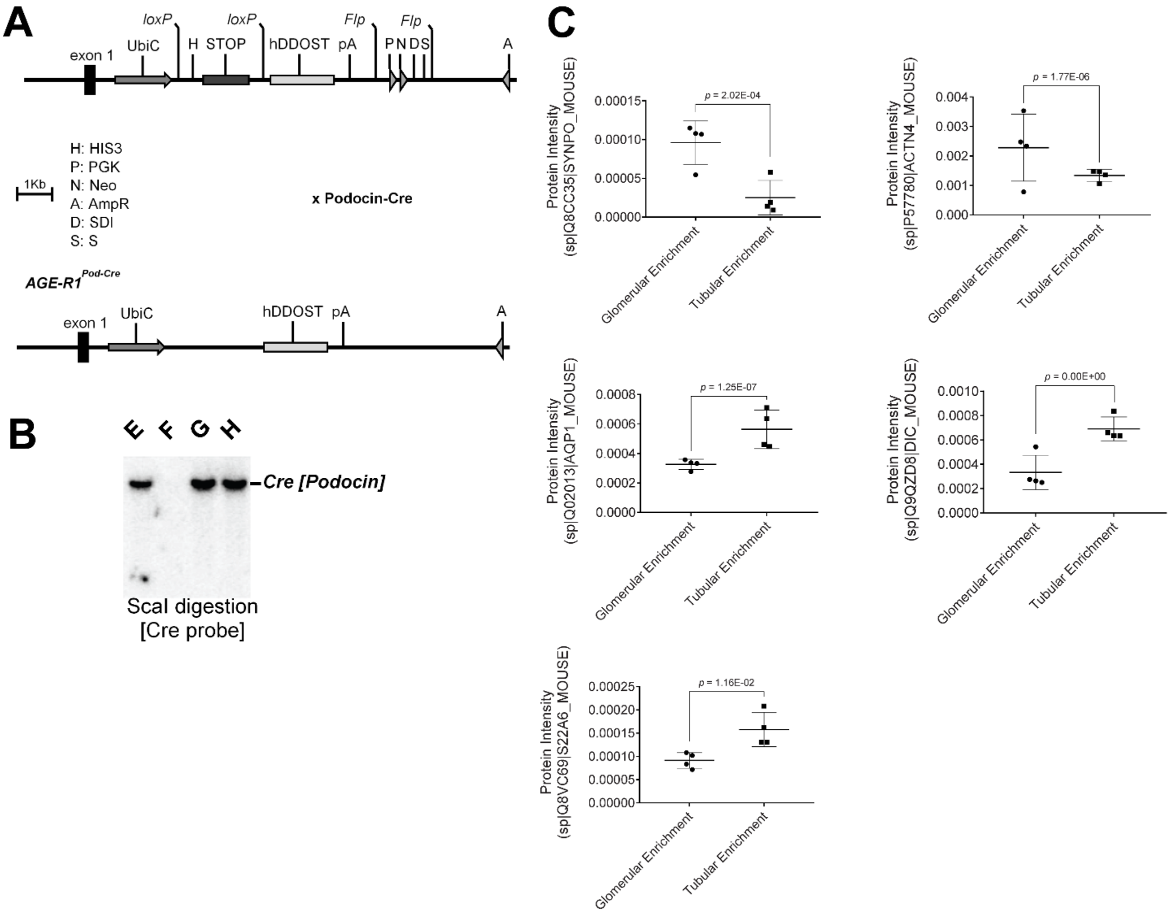

**Supplementary Figure 2.**

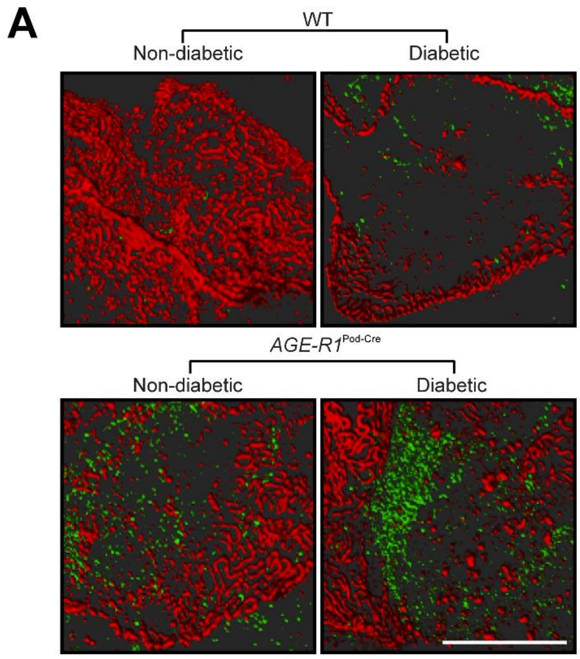

**Supplementary Figure 3.**

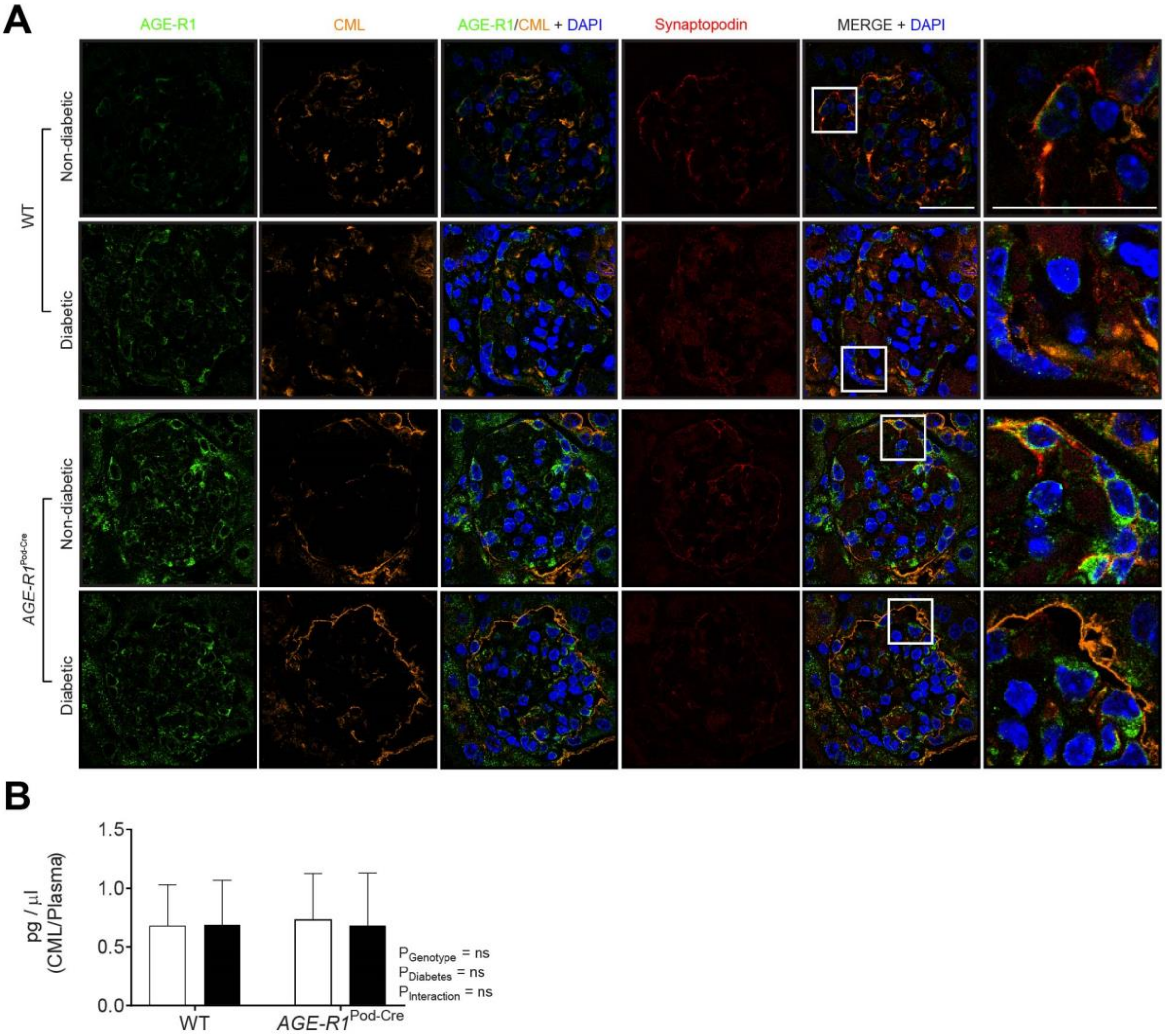
